# Tumour suppressor WT1 and its interacting protein RNA helicase DDX5 regulate the microRNA let7/Igf1r axis in the kidney mesenchyme

**DOI:** 10.1101/2025.01.05.631347

**Authors:** Sourav Dey, Joan Slight, Stuart Aitken, Iwin Joseph, Utkarsh Bhore, Alex vonKriegshiem, Giulia Petrovich, Jocelyn Charlton, Viktoriya Stancheva, Debasis Mohapatra, Abdelkader Essafi, Kathy Pritchard-Jones, Nicholas D Hastie, Ruthrothaselvi Bharathavikru

**Affiliations:** Department of Biological Sciences, Indian Institute of Science Education and Research, Berhampur, Ganjam, Odisha 760010; MRC Human Genetics Unit, Institute of Genetics and Cancer, University of Edinburgh, Western General Hospital, Crewe Road (S), Edinburgh, EH4 2XU; University of Dublin, Ireland; Department of Oncology, Great Ormond Street Hospital for Children, NHS Foundation Trust, Great Ormond Street, London WC1N 3JH, UK; Parala Maharaja Engineering College, Sitalapalli, Berhampur, Odisha, 761003, India; School of Cellular and Molecular Medicine, Faculty of Biomedical Sciences, University of Bristol, University Walk BS8 1TD

**Keywords:** Wilms Tumour, microRNA, DDX5, WT1, IGF

## Abstract

Germline mutations in tumour suppressor, Wilms Tumour 1 (*WT1*) as well as somatic and germline mutations in microRNA processing genes (MIRPG) have been identified in Wilms’ Tumours (WT), where upregulation of IGF2 through genetic and/or epigenetic mechanisms is commonly seen. Whether these different epigenetic and genetic causes converge on common targets is still unclear. WT1 is also involved in RNA binding and regulates the stability of these developmental mRNAs. We now show that WT1 interacts with microRNA let-7 and proteins involved in microRNA processing wherein DDX5 is found to be an important regulator. WT1 absence results in decreased microRNA levels in cell lines and kidney mesenchyme, which is further downregulated in a WT1 and DDX5 double knockdown, ultimately modulating *Igf1r* expression. These findings suggest a plausible mechanism by which *WT1* mutations lead to WT, and converge on microRNA pathway and IGF signalling pathway, which are important contributing factors in the aetiology of WT.

## Introduction

Wilms’ Tumour (WT) is a paediatric kidney disease that may arise from germline mutations in the tumour suppressor genes *WT1* or *WTX*. The incidence of WT is approximately one in every 10,000 (**Charlton et al. 2016**). Germline mutations of *WT1* also lead to other genitourinary abnormalities, and, in some rare cases, congenital diaphragmatic hernia (**Hastie et al. 2017**). Some of the overgrowth syndromes, such as Beckwith–Wiedemann Syndrome (BWS) and Perlman Syndrome, also have an increased risk of Wilms’ Tumour (**Bharathavikru et al. 2018**). The mechanisms contributing to WT are diverse, complex and not completely understood. For example, both *WT1* mutations and *IGF2* upregulation (brought about by genetic and/or epigenetic changes), have now been shown to be present in WT as well as BWS patients. This is also true in the mouse model, where *Wt1* deletion along with activation of Igf2 signalling leads to WT (**Hu et al. 2011**). Other categories of mutations include stabilization of β catenin and p53 mutations (especially in some anaplastic tumours) (**Bardeesy et al. 1994)**. Additional mutations in WT have been identified with patient samples using a targeted exome sequencing approach. Apart from the genes that are involved in kidney development, a new category of mutations in the microRNA processing pathway (**Wegert et al. 2015; Gadd et al. 2017; Ludwig et al. 2016**) has been identified.

Thus, the driver mutations can now be divided into two major categories; nephron differentiation (*WT1, SIX2, METT*), and the microRNA processing gene (MIRPG) pathway (*DICER, DIS3L2, DROSHA, DGCR8*). Few of the most recent studies have also identified predisposing mutations in chromatin modifiers such as *KDM3B* (**9**). The microRNA processing pathway is a highly co-ordinated regulatory programme that regulates RNA levels, thereby affecting protein levels as well. The microRNA pathway consists of the following steps: i) transcription of the primary microRNA (pri-microRNA) from the genome by a Pol-II driven mechanism, ii) followed by the action of the microprocessor complex, composed of DGCR8 and DROSHA, which recognises the stem loop structure and the sequence of the pri-microRNA, which is then cleaved to form the premature microRNA (pre-microRNA), iii) The pre-microRNA is then exported to the cytoplasm by the exportin 5 complex, iv) where the double-stranded pre-microRNA is then recognized by DICER, which along with other proteins, further processes it and forms mature microRNA. The mature microRNA consists of two strands, the guide RNA and the passenger RNA, v) The guide RNA is then loaded onto the RNA induced silencing complex (RISC) complex, that comprises Argonaute (AGO 1/2). The RISC is then capable of silencing gene expression by different mechanisms that include deadenylation, translational repression and degradation mainly by targeting the 3’UTR of the mRNAs (**Winter et al. 2009**) (**Fig. 1A**). Several other RBPs have been shown to interact with these core components of the microRNA processing pathway, modulating the microRNA levels. One such protein is the DEAD box RNA helicase, p68/DDX5 which was initially shown to be associated with the Drosha complex and regulates processing of a subset of primary microRNA. A general role for DDX5 in unwinding of the microRNA let-7 duplex and facilitating RISC complex assembly has highlighted its role in the final step of microRNA targeting to 3’UTR (**Xing et al. 2019**).

**Figure 1:**
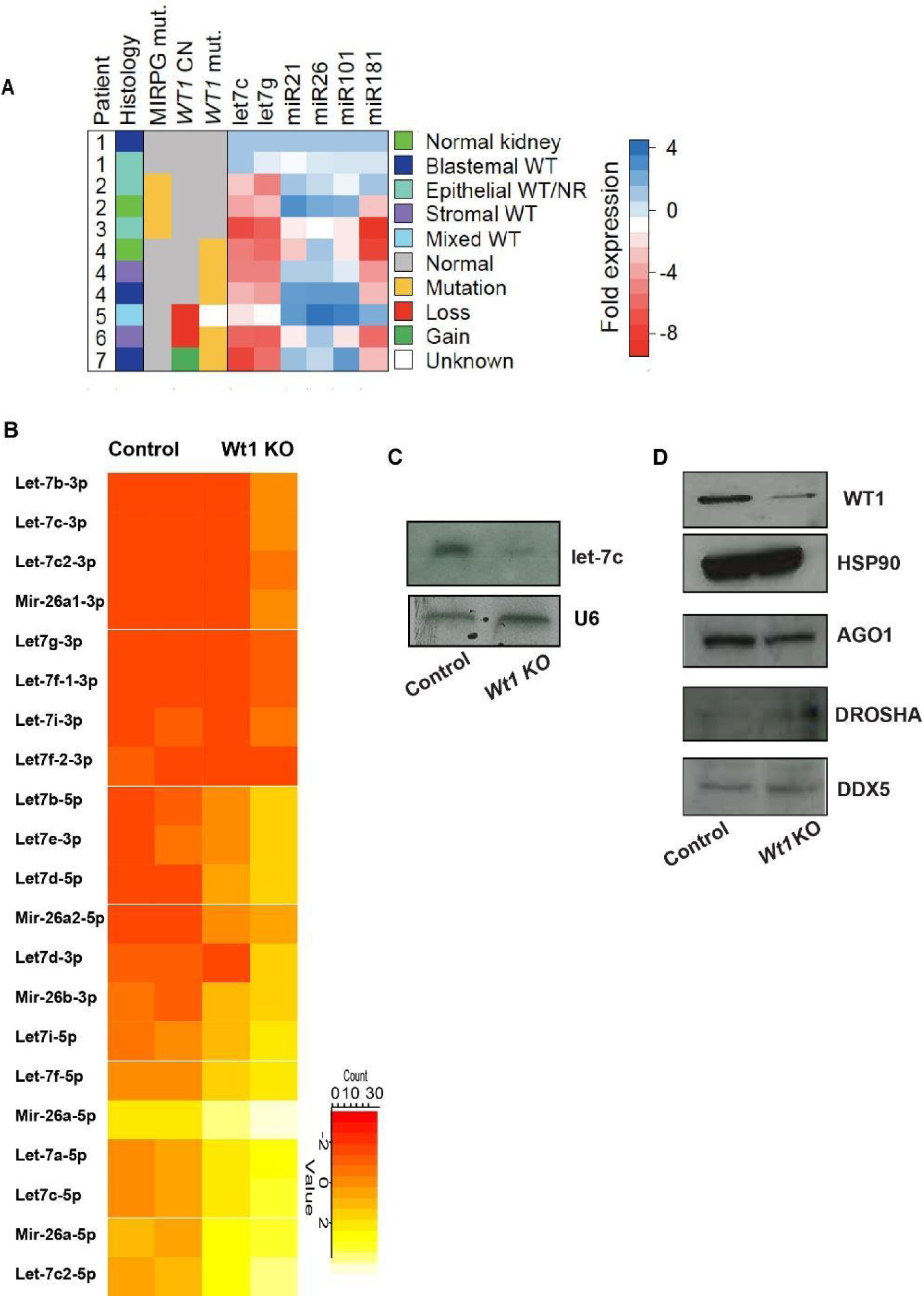
Tumour suppressor WT1 regulates microRNA let-7 expression. **A)** For seven patients, the histology, mutational status for MIRPG, copy number for WT1 (MLPA) and mutational status for WT1 is displayed. MIRPG mutants included DGCR8 (patient 2) and DICER (patient 3). WT1 mutations included 1568C>CT, 458R>R/X, 948_949het_insG (patient 4), Ins CGTAGAC (patient 6) and NM_000378:c.G208A:p.G70S (patient 7). For the microRNAs displayed, the fold change in expression (based on qPCR) is shown, relative to patient sample 1, with normal expression of MIRPG genes and *WT1*. **B)** Heat map representation of differentially regulated microRNAs of the let-7-family, in the 5-day retinoic acid treated ES cells (Wt) compared to the Wt1 deleted ES cells (KO). The heatmap representation ranges from the low expression level of −5 to high expression level of +5, with a median change of 2 in most cases. **C)** Northern blotting analysis of mature let-7c expression compared to U6 loading control in ES cells compared to Wt1 knockout cell line. **D)** Western blotting analysis of ES cells compared to the knockout cell line, showing no observable changes in the levels of the microRNA processing pathway proteins (AGO1, DROSHA, DDX5). HSP90 is used as a loading control. WT1 levels depict the knockout.

The MIRPG mutations identified in WT are mostly somatic, except those in *DICER*, which in some cases were also germline (**Palculict et al. 2016**). The microRNA pathway has also been indirectly linked to WT. Recent studies have shown that the MIRPG-mutation associated with WTs also converges on the IGF signalling pathway through transcriptional mechanisms (**Hunter et al. 2017**). The Perlman overgrowth syndrome, characterized by an increased susceptibility to WT, arises because of mutations in the exoribonuclease *DIS3L2*, which is involved in degradation of oligouridylated transcripts, including microRNA let-7 (**Astuti et al. 2013, Chang et al. 2013**). One of the recent mouse models of WT was created by overexpressing LIN28 in the kidney mesenchyme where WT1 is usually expressed (**Urbach et al, 2014**). The RNA binding protein LIN28 is involved in the homeostasis of the microRNA let-7 pathway by binding to let-7 microRNAs and preventing their translational repression (**Tsialikas et al. 2015, Viswanathan et al. 2008**). Although a global reduction in microRNA levels was observed in the LIN28 WT mouse model study, the introduction of let-7g resulted in tumour regression, suggesting the let-7-LIN28 microRNA pathway to be the one of the underlying cause of WT manifestation and tumour pathogenesis (**Urbach et al, 2014**).

Wilms Tumour 1 (WT1) protein is one of the major tumour suppressor proteins that is associated with WT. The transcription factor role of WT1 has been considered to be responsible for the cellular changes observed in WT. However, we have previously reported that WT1 is also a regulator of RNA stability and that its role in posttranscriptional regulation is equally important in development (**Bharathavikru et al. 2017**) where WT1 interaction with mRNAs was observed. In the same study, we observed WT1 interaction with noncoding RNAs including microRNAs. We decided to investigate if WT1 played a role in the microRNA processing pathway, especially in the let-7 microRNA pathway, since the let-7-LIN28 pathway has been shown to be associated with WT.

In this study, we show that in WTs, there is a decrease in the mature microRNA let-7 levels in patient samples with WT1 mutation or loss. In the absence of WT1, microRNA let-7 expression is downregulated in the ES cells and mesonephric cells, which is regained by a transient induction of WT1. The most significant category of microRNAs that responds to the levels of WT1 is the let-7 family. An unbiased protein protein interaction (PPI) study was performed for endogenous WT1 which revealed the enrichment of RNA processing pathways, including the microRNA processing pathway. Detailed analysis of these PPI networks highlighted DDX5 as one of the regulatory nodes in the pathway. WT1, thus interacts with the microRNA processing pathway proteins and modulates microRNA expression. WT1 and DDX5 interaction was found to be a direct interaction based on endogenous Immunopulldowns as well as Chemical Induced Dimerization-Fluorescence Resonance Energy Transfer (CID-FRET) based analysis. A combined knockdown of WT1 and DDX5 showed additive downregulation of microRNA let-7 expression. Further analysis of let-7 targets shows a significant upregulation of *Igf1r* expression that impinges on the IGF signalling pathway. The WT1-let-7-*Igf1r* axis was also validated by using an inhibitor against microRNA let-7 which sequesters the existing microRNA. In Wt1 kd cells, the level of *Igf1r* was further increased upon miR let-7 inhibitor treatment. The microRNA processing defect is also observed in kidney mesenchyme deleted for *Wt1* using a Nestin-cre *Wt1* conditional mouse model. All these observations suggest that the dysregulation of the microRNA let-7 pathway and the downstream *Igf1r* modulation is one of the contributing causes of WT.

## Results

### Tumour suppressor, WT1 regulates microRNA let-7 expression

*WT1* is frequently mutated in paediatric kidney cancer, WT, and several studies have shown that microRNA-processing pathway genes (MIRPG) are also mutated with an associated decrease in mature microRNA levels (**Wegert et al. 2015, Gadd et al. 2017, Ludwig et al. 2016, Torrezan et al. 2014, Rakheja et al. 2014**). To assess whether *WT1* mutations may also impact microRNA expression levels, we performed a small-scale analysis with seven patient samples. The expression levels of six microRNAs were assessed by qPCR in a representative panel of WT and normal kidney samples with *WT1* mutations or an aberrant gene copy number, as well as samples with MIRPG mutations (*DICER* or *DGCR8*) as positive controls (**Fig. 1A**). In comparison to unmutated WTs, MIRPG samples displayed altered microRNA expression levels, including downregulation of let-7c, let-7g and miR181 (**Fig. 1A**). Supporting the role of WT1 in microRNA processing, samples with *WT1* mutation showed remarkably similar microRNA expression levels to MIRPG mutant samples (**Fig. 1A**). Hence, we decided to investigate if Wilms Tumour suppressor protein, WT1 also modulates microRNA let-7 pathway by analysing expression in *Wt1* deleted knockout (KO) embryonic stem (ES) cells, using a microRNA profiling approach (**Fig. S1A, S1B**). The let-7 family of microRNAs were found to be differentially regulated corresponding to the presence of WT1. In the absence of WT1, all members of the let-7 family showed decreased expression by at least 2-fold in comparison to the control (**Fig. 1B**). Northern blotting analysis of cells where WT1 is depleted (*Wt1* knockout embryonic stem (ES) cells or *Wt1* knockdown mesonephric cells (M15), showed a downregulation of mature microRNA let-7c levels (**Fig. 1C, S1C resp.**). A qRT PCR analysis of the premicroRNA let7 levels in Wt1 KO ES cells and Wt1 Kd M15 cells showed an upregulation (**Fig S1D, Fig S1E resp**.) suggesting that, in the absence of WT1, there is a defect in the microRNA processing pathway.

Earlier studies have shown defective microRNA processing when MIRPG is deleted (**Lee et al. 2004, Yermalovich et al. 2019**). Hence, to assess whether there was any change in the expression of the microRNA pathway proteins, western blotting analysis of DROSHA, DDX5 and AGO proteins was performed. We did not see any significant changes of these proteins, corresponding to the presence or absence of WT1 both in ES E14 cells and M15 cells respectively (**Fig. 1D, S1F**). Steady-state levels of let-7 microRNAs are regulated by the RNA binding protein LIN28 **(Tsialikas et al. 2015, Viswanathan et al. 2008**). It has also been shown that the let-7 microRNA is an important regulator of timing during nephrogenesis (**Yermalovich et al 2019**). In ES cells, WT1 expression increases during a 5-day retinoic acid (RA) time course (**Bharathavikru et al. 2019**, **Spraggon et al. 2007**). During the same time period of differentiation, let-7 levels also increase (**Fig. S1G**) with a concomitant decrease in LIN28 expression (**Fig. S1H,** ES panel, day 5 of RA treatment). Therefore, we decided to investigate whether LIN28 plays a role in WT1-mediated microRNA processing. As expected, Wt1 levels increase across the 5 days of RA treatment in the wildtype cells, whereas the KO ES cells do not respond in a similar manner (**Fig. S1H**). Lin28 levels, however, showed minimal differences between wildtype and knockout cells across the same time course (**Fig. S1H**). Thus, the observed defect in the microRNA processing, in the absence of WT1, is not due to any changes in the expression of Lin28 or the microRNA processing pathway proteins.

To ascertain whether WT1 directly regulates microRNA levels, we decided to use a gain of function approach. Knockout ES cells were transiently transfected with a plasmid expressing the +exon5/+KTS isoform of WT1 (**Fig. S2A**). In the plasmid, the expression of WT1 is under the influence of doxycycline (dox) regulation, hence, dox treatment was used to titrate the levels of WT1. Induction of WT1 was confirmed by imaging and western blotting (**Fig. S2B, S2C**). Accordingly, with dox induction, there was an increase in the let-7 microRNA levels observed in comparison to the corresponding no dox treatment controls. To visualize the overall pattern of microRNA expression changes, we performed microRNA sequencing and compared the expression of 637 detectable microRNAs across four samples with varying expression levels of WT1: knockout ES cells without dox (KU), dox 12 hours (D12), dox 24 hours (D24) and ES cells treated with retinoic acid for 5 days (ERA) (**Fig. 2A**). We correlated the expression of each microRNA with WT1 expression across the four samples and observed a highly correlated cluster of microRNAs that included let-7 microRNAs (**Fig. 2A**). Indeed, the expression of 9 detectable mature let-7 transcripts appeared to increase with higher WT1 levels (i.e. from Ku to ERA, (**Fig. 2B)**. It can be concluded that the transient induction of WT1 is sufficient to rescue the microRNA levels, and especially of microRNA let-7, in Wt1 knockout ES cells. This clearly shows a regulatory role of WT1 in the let-7 pathway.

**Figure 2:**
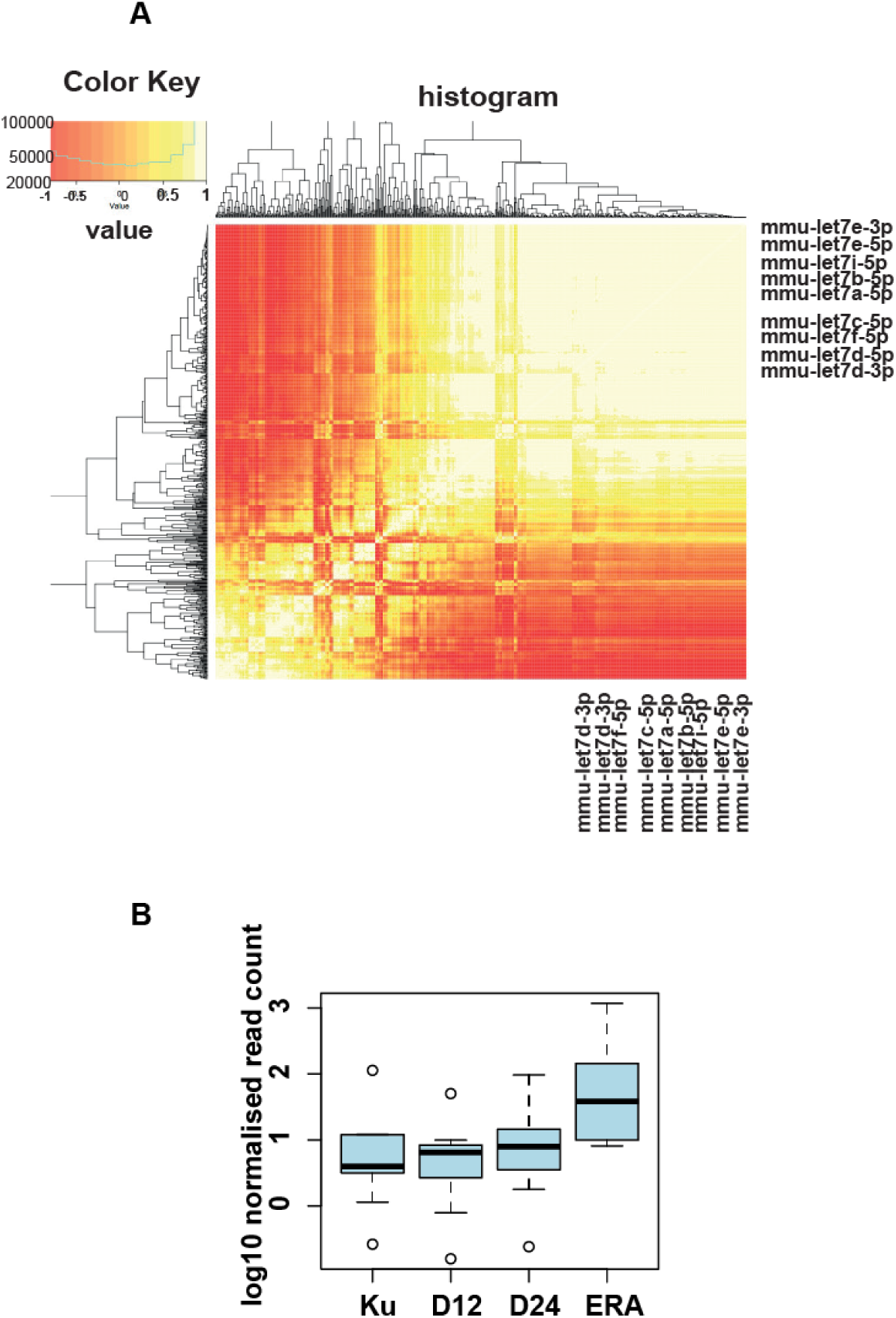
Tumour suppressor WT1 regulates microRNA let-7 expression through transient induction. **A)** Heatmap of pairwise correlations of WT1 expression in four conditions (Ku, D12, D24 and ERA, see main text) to 637 expressed mature microRNA. A large proportion of microRNA have highly correlated expression patterns (clustered top right) and this population includes let-7s. **B)** Boxplot representation of let-7 levels for 9 mature let-7 microRNA (let-7a,b,c,d,e,f,i 5’ transcripts and let-7d,e 3’ transcripts) in samples across uninduced and WT1 transfected KO ES cells, induced with doxycycline for 12 hours and 24 hours and wild type ES cells treated with RA.

### Transcription factor and RNA binding protein, WT1 interacts with microRNAs

The global analysis of WT1-interacting RNA (**Fig. S3A**), by FLASH (FormaLdehyde-Assisted crosslinking and Sequencing of Hybrids) and RNA immunoprecipitation (RIP), was done in mesonephric M15 cells (**19**). This showed different noncoding and small RNA subtypes, of which microRNA was one of the common enriched categories (**Fig. 3A, 3B**). The comparison of WT1-interacting microRNA to the input control identified by FLASH is depicted in (**Fig. 3B)**, which shows a characteristic bimodal pattern of enrichment. The interaction with microRNAs was confirmed by northern blotting using probes specific for the mature microRNAs let-7c, let-7g and U6 snRNA as the control (**Fig. S3B, S3C**). This indicates that WT1-microRNA let-7 interaction plays an important role in the microRNA let-7 pathway, resulting in altered let-7 expression in the absence of WT1. We have previously identified the RNA binding motif for WT1, enriched mostly in the 3’ UTRs of mRNAs and ncRNAs including microRNAs as shown in (**Fig. 3A)**. However, detailed analysis was restricted to the 3’ UTR of the mRNAs corresponding to developmental pathways. A comparison of the microRNA-mRNA hybrids obtained from FLASH and the mRNA expression from transcriptome analysis reported earlier, shows very little overlap between the targets (**Fig. 3C, Table S1**) suggesting that WT1 mediated microRNA regulation might not be only through microRNA interaction but through other regulatory mechanisms.

**Figure 3:**
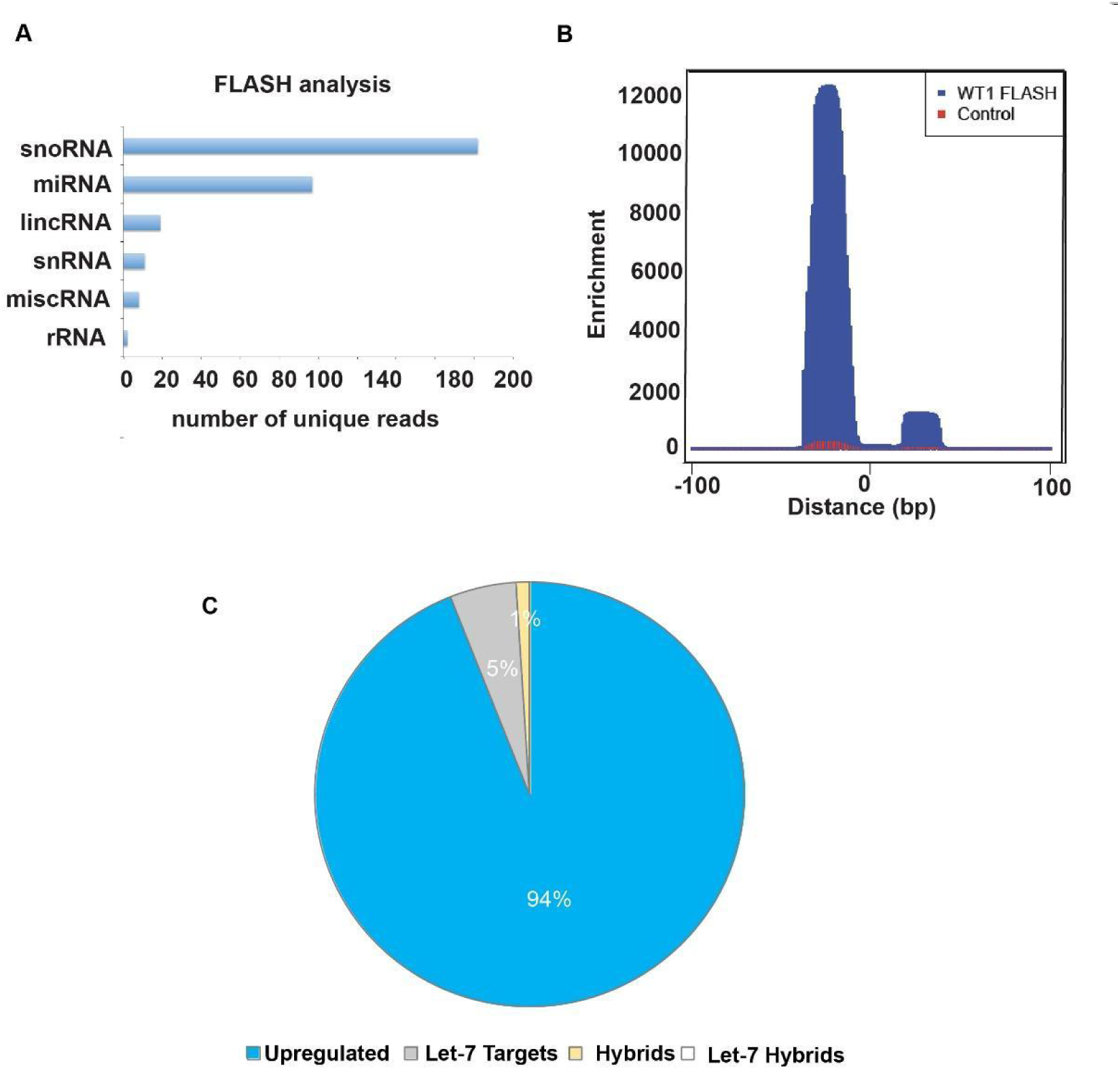
Transcription factor and RNA binding protein, WT1 interacts with microRNAs. **A)** Graphical representation of the percentage of WT1 interacting noncoding biotypes; lincRNA, microRNA, rRNA, snRNA, snoRNA, misc RNA in FLASH (FormaLdehyde-Assisted cross linking and Sequencing of Hybrids) analysis of mesonephric cell line, M15. **B)** Read pile-up plot of microRNAs in M15 WT1 FLASH analysis compared to input control. The x-axis represents the 200-base pair window, centered in the middle of each microRNA. **C)** Pie chart representing the overlap between the microRNA-mRNA hybrids FLASH data and the transcriptome data for M15 cells compared to the *Wt1* kd cells.

### WT1 interacts with microRNA processing pathway proteins including the RNA Helicase DDX5

In order to gain insights into whether WT1 regulates microRNA processing through protein-protein interactions, WT1 Immunopulldowns (IPs) was performed as described in **Bharathavikru et al. 2016,** and subjected to a mass spectrometric analysis (**Fig. S4A**). The identified proteins were analyzed and ranked according to the intensity values. This list was analyzed using GOrilla and ShinyGO tools (http://bioinformatics.sdstate.edu/go77/, https://cbl-gorilla.cs.technion.ac.il) to understand the relative enrichment of specific pathways/gene ontology. Majority of the proteins were found to be associated with RNA processing, and Translation related pathways (**Fig. 4A**). We generated an interaction plot by Shinygo of all the biological processes, where three different clusters were observed, of which gene ontology associated with RNA processing and regulation processes was one of the major clusters (**Fig. S4B)**. This observation strengthens the role of WT1 in various steps of RNA processing. String database analysis of the immunoprecipitated proteins revealed three clusters of interacting protein networks. Of these three different clusters, the green cluster represents proteins involved in RNA processing (**Fig. 4B**). This cluster was further analyzed in detail to understand the main regulatory nodes with Cytoscape, wherein, we identified DDX5, as the central node of interaction (**Fig. 4B, inset**). In another related study using M15 genome edited cells that overexpress the +KTS isoform, the endogenous WT1 IP and mass spectrometry data was subjected to a graph theory based ranking analysis. In this study as well, DDX5 was identified as one of the major interacting proteins with a high betweenness centrality score that depicts the interaction probability of WT1 and DDX5 (**Fig. 4C, Table S3**).

**Figure 4:**
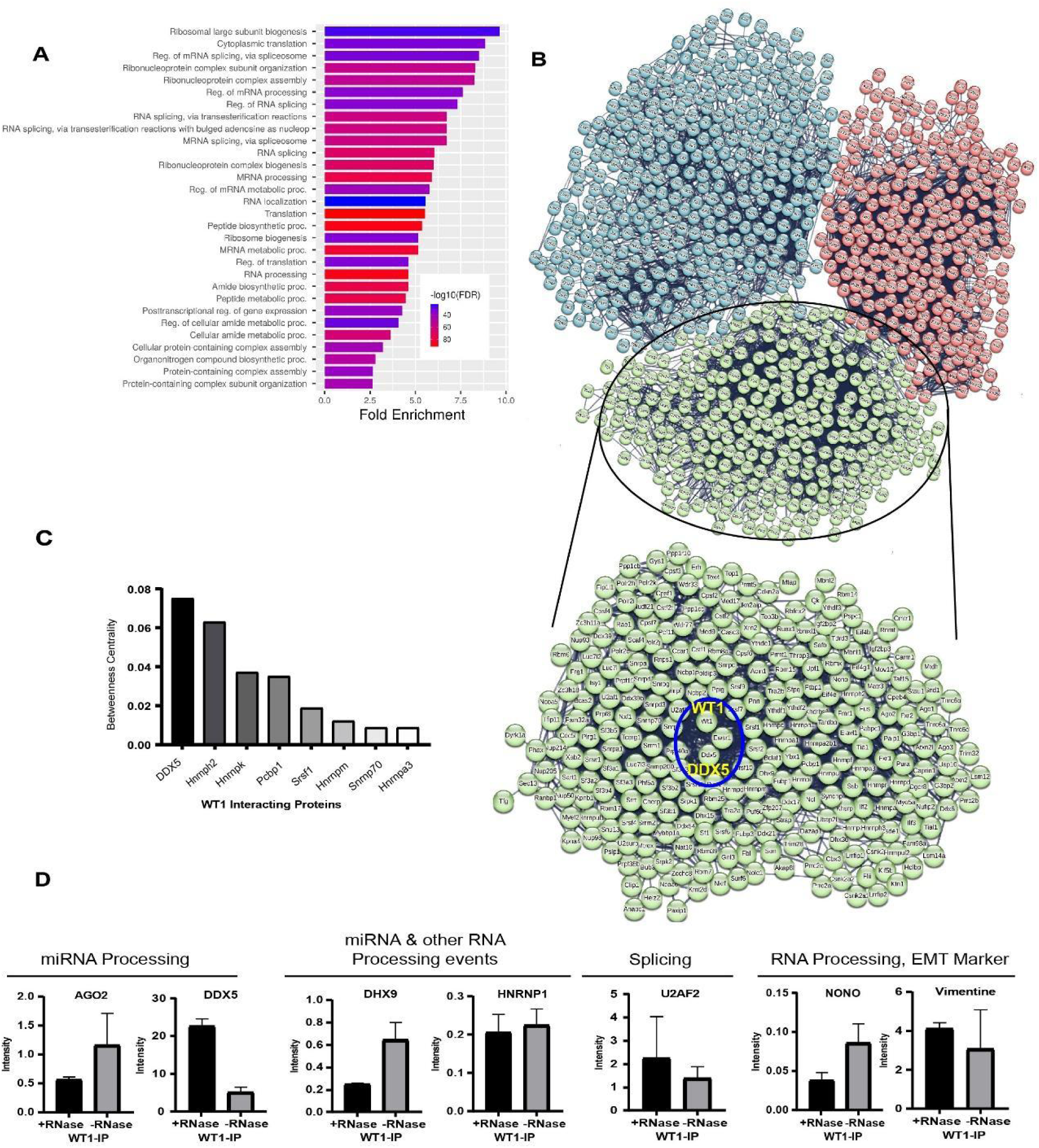
DDX5 as a Central node of WT1 interacting proteomes. **A)** Top 1000 genes from the Mass Spectrometry dataset of WT1-IP samples were used for Gene Ontology analysis by Gorilla and Shinygo, with a 0.05 FDR cutoff, biological processes are represented by a color-coded bar graphic that reveals RNA processing and translation regulation as processes that are highly enriched. **B)** The listed proteins were analyzed using the STRING database to create a protein network. This analysis identified three distinct functional clusters formed by the immunoprecipitated proteins, Ribosome biogenesis (Red), Amino acid metabolism and translation (Blue) and RNA processing and Translation regulation (Green). The cluster depicting the RNA processing biological processes is presented as an inset, which shows DDX5 as the central node and key connector to WT1. **C)** Applied graph theory-based analysis of WT1 PPI represented on the basis of ranking of betweenness centrality score **D)** Relative enrichment of representative WT1 interacting proteins (microRNA processing, other RNA processing, Splicing, EMT), identified by Mass Spec analysis of WT1-IP are represented graphically. Enrichment value is calculated over IgG Control pulldown.

From the interactome, we categorized the interacting proteins according to their biological activity (**Xing et al. 2019, Rodriguez et al. 2023**). A graphical representation of few of the interacting proteins that were identified in the above analysis in the presence or absence of RNase treatment shows a differential pattern of enrichment for AGO2, DDX5, DHX9, HNRNPA1, U2AF2, NONO & Vimentin. Notably, DDX5 shows an RNA independent interaction with WT1, suggesting that it could be a direct protein-protein interaction (**Fig. 4D**). Upon ranking of all the RNA processing events associated with DDX5 and WT1 interaction, we identified the microRNA mediated gene silencing by translation inhibition as a significantly enriched term (**Fig. S4C**).

We performed immuno pulldowns to validate the microRNA processing pathway proteins. It has been previously shown that WT1 interacts with the microRNA pathway components DICER and AGO2 (**Apka et al. 2016**). WT1 interaction with AGO1, AGO2 and DDX5 was confirmed in the mesonephric cells through WT1 IPs (**Fig. 5A**). In order to confirm whether WT1-DDX5 is a direct interaction we resorted to a Chemically Induced Dimerization (CID)-Fluorescence Resonance Energy Transfer (FRET) based interaction assay. CID, is a chemically regulated assay to study protein interaction, and has been earlier used to study the interaction of the proteasome adapter to the target proteins (**Wilmington et al. 2016**) whereas FRET based PPI analysis has been used in multiple studies to understand protein-protein interaction in live cells (**Banning et al. 2010, He et al. 2003**). We have generated the WTI-DDX5 CID system by creating fusion proteins in the fluorescence vector background to obtain cells that express DDX5-CFP-FKBP and WT1-YFP-FRB expressing cells. FKBP and FRB both contain Rapamycin binding pockets, and can form a dimer only in the presence of the chemical, Rapamycin. Before initiating the CID FRET based analysis, a confocal imaging was performed to assess the colocalization of WT1 (YFP tagged) and DDX5 (CFP tagged), which showed a colocalization rate of 88.92 (**Fig. 5B, Table S4**). Various concentrations of Rapamycin were tested to optimize the CID, measured as a FRET readout using Flow cytometry calculated as the change in FRET ratio (**Fig. S5A**).

**Figure 5:**
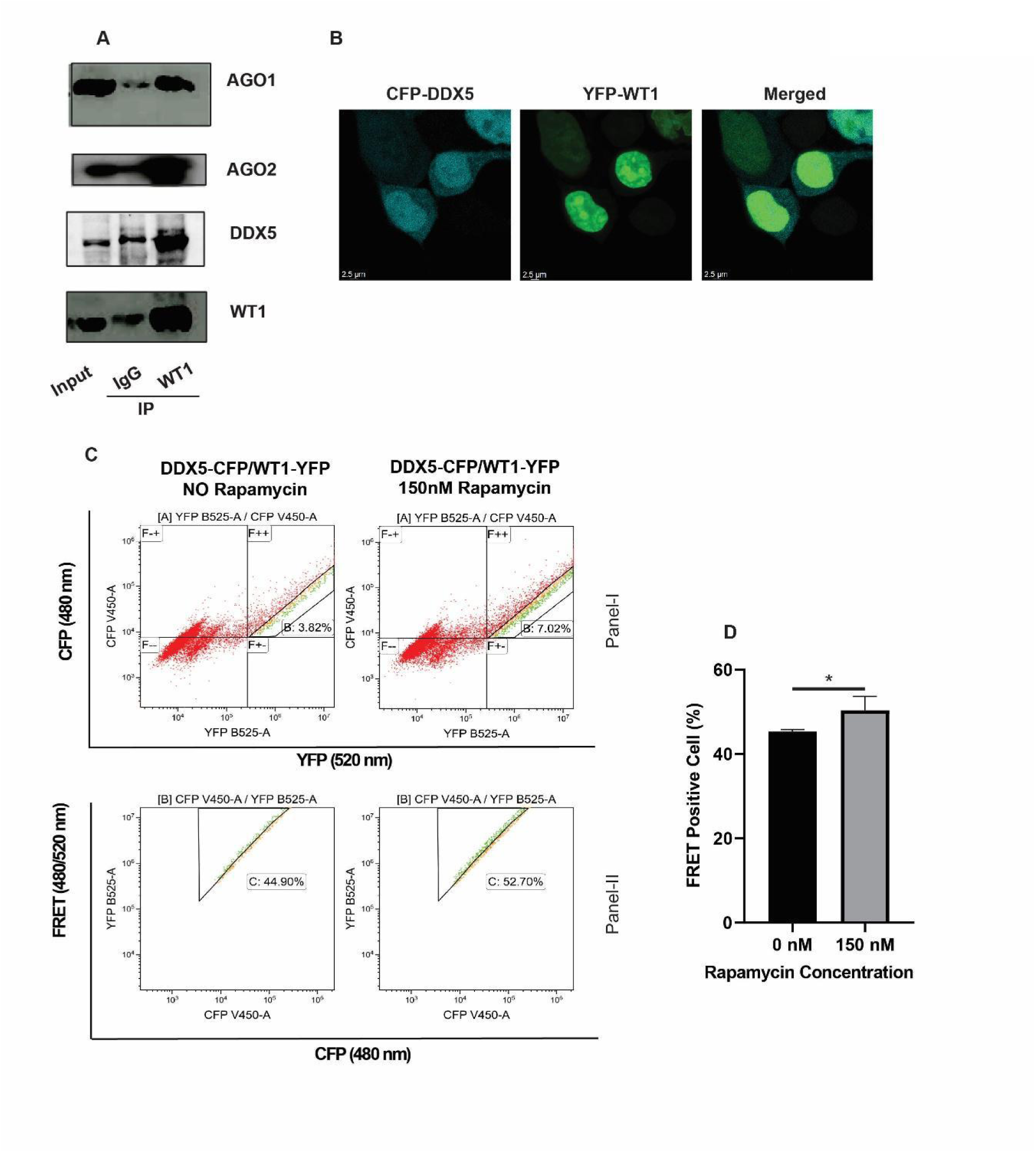
WT1 interacts with RNA Helicase DDX5. **A**) Immuno-pull down by WT1 antibody and IgG antibody as a control to validate WT1 interacting proteins (AGO1/2, DDX5) along with 10% input. **B**) Confocal imaging of CFP-DDX5 and YFP-WT1 shows co-Localization of WT1 and DDX5 **C**) Chemically Induced Dimerization and FACS based FRET analysis performed to study the Interaction strength of DDX5 and WT1. Rapamycin (150 nM) was treated to the CFP-DDX5 & YFP-WT1 (Right) transfected cells, and FRET positive cells were recorded on CFP/YFP (V450/B525) channel. **D**) Graphical representation of the percent of FRET positive cells for both with (right) or without (left) chemical treatment that were analysed by KALUZA Software.

The FRET ratio was calculated by excluding the double positive cells that arise from cotransfection of CFP and YFP only then analyzed the true FRET positive cells in panel-II (**Fig. S5B**). False positive cells that were excited by 430 nm laser were gated out and then the double positive cells were recorded in Panel-I. FRET positive cells were recorded with various concentrations of rapamycin across time and represented graphically in (**Fig. S5C)**. With the optimized conditions, Interaction between DDX5 and WT1 observed by a flow cytometry-based FRET (**Fig. 5C**). FRET positive cells with or without rapamycin treatment were analyzed and represented graphically (**Fig. 5D**). A similar number of FRET positive cells, with very minimal variation on rapamycin addition was observed, indicating that the interaction of WT1 and DDX5 is independent of the CID system, and hence a direct protein-protein interaction. As a control, WTAP, which is an established WT1 interacting protein, was also investigated for its protein-protein interaction through CID FRET analysis as shown in (**Fig S5D)**. The role of DDX5 in microRNA processing has been observed earlier. In this study, we show that a tissue specific developmental regulator, WT1, through its direct interaction with DDX5 can regulate microRNA expression.

### WT1 and DDX5 regulate microRNA let-7 processing

To understand the potential role of WT1 and DDX5, together in regulating microRNA expression, we created genome-edited lines of DDX5 in the *Wt1* kd M15 cells (double kd). M15 cells that were subjected to genome editing to create a DDX5 kd (single kd) were used as controls. Two gRNA targets, sg1 and sg2 were selected to create the DDX5 knock down in both M15 and MW cells (MW-WT1 knockdown M15 cells), which was confirmed by western blotting (**Fig. 6A. Panel I, and Panel II, single kd and double kd resp**). Fold expression of DDX5 in both cells with respect to empty vector treated control cells was assessed by ImageJ (**Fig. S6A**), which shows an almost complete knockout with sg1 in single kd and approximately 50% reduction in the double knockdown. qPCR analysis of mature let-7c microRNA expression status in both DDX5 kd cell lines revealed a downregulation of about 40% let-7 expression in single kd cells. The double knockdown cells showed a significant additive downregulation (about 60-65%) of let-7 expression (**Fig. 6B, Panel I (single kd), Panel II (double kd) and Panel III (combined representation)**), thus leading to the conclusion that WT1 and DDX5 cooperatively regulate microRNA let-7 expression in the mesonephric cells.

**Figure 6:**
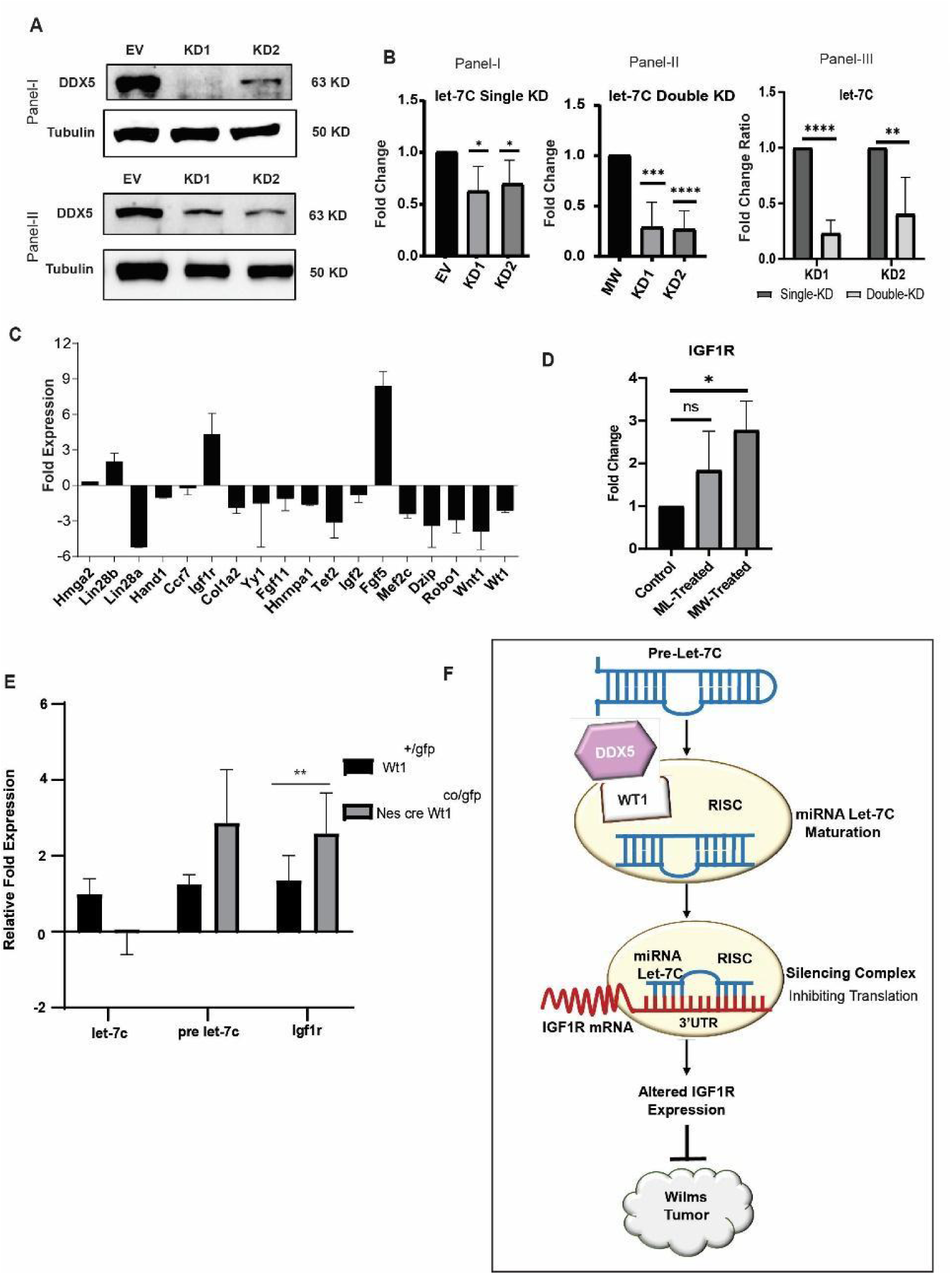
WT1 regulates microRNA let 7 processing through DDX5. **A)** Western blotting images of DDX5 knockdown in M15 (Panel I) and WT1 knockdown M15 cells (MW), (Panel II) respectively. **B)** Expression of microRNA Let-7c was studied by qPCR from both single (Panel I) and double DDX5 knockdown (Panel II) cells and a combined representation in Panel III. Error bars represent the mean and SD, p<0.05. **C)** qPCR analysis of TARGETSCAN-predicted let-7 targets, in the presence (*lacz kd*) or absence of WT1 (*Wt1 kd*), represented as fold change in expression, normalized to 18s RNA expression. **D)** *Igf1r* expression studied by qPCR analysis upon let-7 inhibitor treatment in M15, control knockdown and Wt1 knockdown cells. Untreated M15 used as control. Error bars represent mean and SD. p<0.05. **E)** qPCR analysis of pre-microRNA let-7c, microRNA let-7c, *Igf1r* in *Nestin Cre* WT1 conditional mutants, shows downregulation of microRNA let-7 expression and upregulation of *Igf1r* compared to wild type WT1 expressing cells. **F)** Schematic to explain the role of WT1 in the let-7-*Igf1r* axis in the kidney mesenchyme. WT1 regulates Igfbp5 mRNA through regulating RNA stability. WT1 also regulates let-7 microRNA processing through interaction with DDX5 and thus influences *Igf1r* expression. In the absence of WT1, the Igfbp5 mRNA is less stable and microRNA let-7 levels are downregulated, correspondingly resulting in increased *Igf1r* expression, thus resulting in increased IGF signalling with potential implications in Wilms’ Tumour manifestation.

### The WT1-microRNA let-7*-Igf1r* axis in the kidney mesenchyme

The physiological relevance of the regulation of let-7 microRNA levels will be evident through the expression of their mRNA targets. Since let-7 microRNA levels are downregulated in the absence of WT1, upregulation of let-7 target mRNAs would be expected. To investigate this, we analyzed the levels of a representative subset of TargetScan predicted let-7 targets in Wt1 knockdown M15 mesonephric cells. Two of the predicted let-7 targets, Lin28 and, more strikingly, *Fgf5 and Igf1r* showed significant upregulation upon Wt1 knockdown (**Fig. 6C**), whereas the other predicted targets were either unaffected or downregulated. The likely explanation for this downregulation is that most of these are validated WT1 transcriptional targets (eg: *Col1a1, Igf2*), positively regulated by WT1 (**Motamedi et al. 2014**), as well as other downstream pathways regulated by WT1 transcriptional targets. We have previously shown that WT1 stabilizes mRNAs (**Bharathavikru et al. 2017**). Many of the let-7 targets did not show any appreciable 3’ UTR interaction, nor change in expression with WT1. However, *Igf1r*, which does not interact with WT1, showed an upregulation in expression in the absence of WT1 (**Fig. 6C**). In order to confirm the WT1-let-7-*Igf1r* axis, we treated the cells with the inhibitor of let-7C transiently (**Fig. S6B, S6C**), for 48 hours following which *Igf1r* expression was analyzed by qPCR. In Wt1 knockdown M15 cells, upon microRNA let-7 inhibitor treatment, a further upregulation of *Igf1r* expression over the control cells was observed (**Fig. 6D**), thus strengthening the observation of Wt1 regulating the let-7 expression and thereby modulating *Igf1r* levels.

WT1-mediated regulation of the microRNA let-7 pathway was observed in two different cell lines expressing WT1. To confirm whether this was relevant in a physiological context, Wt1 was deleted in the kidney mesenchyme using the Nestin-cre Wt1 conditional mouse line and the Wt1 GFP knockin model (**Essafi et al. 2011**). Embryonic day E13.5, metanephric mesenchyme cells from gfp control (GFP positive, *Wt1*^co/gfp^) and Wt1 deficient (GFP positive, *Nes cre Wt1*^co/gfp^) genotypes were sorted by FACS to isolate the populations of GFP positive cells. RNA purified from the control and mutants was assessed for microRNA levels. In the mutants, mature let-7 microRNA expression was downregulated whereas, pre-mature microRNA let-7 was upregulated consistent with the observation in cell lines (**Fig. 6E**).

Correspondingly, in Wt1-deleted kidney mesenchyme, the *Igf1r* levels show significant upregulation as compared to the controls (**Fig. 6E**), suggesting that the WT1-let-7-*Igf1r* axis is functional in the tissue context. All the observations suggest that WT1 mediated regulation of microRNA let-7 expression is relevant both in the context of mouse nephrogenesis as well as in WT. Thus, it can be concluded that WT1 interacts with and facilitates the processing of a subset of microRNAs (let-7 family) through PPI interaction involving DDX5, leading to the modulation of the IGF signalling pathway (**Fig. 6F**) through *Igf1r* levels.

## Discussion

In this study, we show that the tumour suppressor protein, WT1 interacts with microRNAs (let7) as well as with the microRNA pathway proteins (DDX5) and thus influences the microRNA processing and the downstream regulation of the target gene Igf1r in kidney mesenchyme.

In the cells that express WT1, we observe that WT1 mutations as well as the MIRPG mutations, all lead to a decrease in the let-7 levels, which would ultimately affect the nephron progenitor differentiation. The RBP LIN28, is a significant protein in the microRNA let-7 pathway. The let-7 microRNA is regulated by the RBP, LIN28, which is essential for maintaining homeostasis. Lin 28 overexpression has been correlated to tumorigenesis (**Viswanathan et al. 2009**). However, we observe minimal changes in the LIN28 levels, although there is a marked decrease in the let-7 levels. In agreement with this, is the recent finding that the lin28-let-7 axis and the timing of let-7 expression regulates nephrogenesis (**Yermalovich et al. 2019**). WT is composed of heterogeneous cell populations that are part of the nephrogenesis pathway, and it has been hypothesized that there are cancer stem cells (CSC) that lead to the manifestation of WT (**Shurkrun et al. 2014**). More recently, CSCs in WT have been identified using SIX2 progenitor cells as markers (**Petrosyan et al, 2023**). In this context, the microRNA pathway, especially the lin28-let-7 axis, could provide significant insights into the causative mechanisms and pathways that underlie WT. Thus, perturbation of the microRNA let-7-lin28 pathway seems to be one of the key pathways that result in WT.

We observe that WT1 interacts with and regulates microRNAs, especially the let-7 family. This is mediated through the direct interaction with the RNA helicase DDX5. WT1 interacts with the microRNAs. This interaction was of a bimodal pattern, suggesting a possible interaction of WT1 with the stem region of the stem-loop structure. WT1 has already been shown to interact with 3’ UTR of mRNAs especially with secondary structures, stabilizing them (**Bharathavikru et al. 2017**), a significant proportion of these mRNAs regulate developmental processes. However, WT1 regulates gene expression through transcriptional and post transcriptional mechanisms and hence majority of let-7 targets when examined in the absence of Wt1 (**Fig. 6C**), show downregulation of expression. The global sequencing experiment revealed different categories of correlation with microRNA and WT1 expression. Within these correlation clusters, it is also possible that there are microRNAs transcriptionally regulated by WT1, and further investigations are required to identify them. This is further supported by the presence of microRNA-mRNA hybrids in the FLASH analysis, which did not show any change in expression; only a few were downregulated in expression (**Table S1, Fig. 3C**). WT1 as a transcription factor is known to activate and repress gene expression (**2**). Further studies on WT1-RNA interaction could possibly highlight such a dual regulatory role in the RNA context as well. The motif analysis shows a preference of WT1 mediated regulation to microRNAs with the UGUA sequence that is found in the let 7 family of microRNAs (**Fig S6D**). The other microRNA families such as microRNA181 which have similar motifs might also be regulated by WT1. Interestingly, the UGUA motif has been shown to be important for CstF recognition (**Hwang et al. 2016**) and SFPQ (**Bottini et al. 2017**).

In this context, the WT1 PPI analysis becomes significant, since WT1 seems to be part of several RNA processing pathways. Considering that WT1 is a tissue specific developmental regulator with restricted spatiotemporal expression, these protein complexes might be crucial in shaping the developmental programme. We have characterized one such interaction, between WT1 and DDX5. The CID FRET based analysis provides evidence for a direct protein-protein interaction. The RNA helicase association with the RISC complex protein Ago2 has already been shown (**Xing et al. 2019**). In this study, we have identified the WT1-Ago interaction to be another PPI in the microRNA pathway, leading us to speculate the existence of a ternary complex involving WT1-AGO-DDX5, which needs to be evaluated structurally.

WT1-mediated let-7 microRNA regulation results in downstream IGF1R upregulation with potential implications in WT aetiology. There are multiple events that lead to WT, including the perturbation of the IGF2 signalling pathway (**Bharathavikru et al. 2018**). This occurs through genetic and/or epigenetic mechanisms including paternal disomy or loss of imprinting of the maternal allele. Recently, it has been shown that microRNA dysregulation can also relate to PLAG1 overexpression that leads to IGF2 upregulation, thus contributing to WT (**Chen et al. 2018**). Additionally, DIS3L2 has also been shown to be involved in the tumour biology by upregulating IGF2 expression (**Hunter et al. 2018**). In agreement with this augmented IGF expression, we observe changes in *Igf1r* levels as a downstream event of let-7 downregulation. WT1 regulates the RNA stability of IGFBPs, including IGFBP5. Both IGFBPs and IGF receptors bind to the growth hormone IGF: IGFBPs sequester IGF and the IGF-IGF receptor interaction potentiates the signalling pathway (**Kim et al. 2009, Chao et al. 2008**). In the absence of WT1, IGFBPs are downregulated (**Bharathavikru et al. 2017**) while *Igf1r* is upregulated, suggesting an increase in the IGF2 signalling events (**Fig. 6F**). Previously, it has been shown that in WT, genetic and/or epigenetic mechanisms lead to increased IGF2 expression, and IGF1R is also upregulated in WT through copy number gain and linked to the possibility of relapse (**Natrajan et al. 2006**). WT1-let-7-*Igf1r* also seems to be an important perturbed node that enhances the IGF signalling pathway and contributes towards the manifestation and pathology of WT. We analyzed the reported miRNA let7 expression data in WT samples that is available on CBioportal for evidence to support the WT-let-7-*Igf1r* axis in patient samples. There was a weak correlation observed between WT1 and let-7 as well as WT1 and Igf1r (**Fig S6E and S6F**). The extent of the involvement of the WT1-let-7 pathway in WT, needs to be addressed in a separate study, involving a larger cohort of patient samples with the entire spectrum of MIRPG mutations.

The lin28-let-7-IGF pathway is a well-established regulatory node in glucose metabolism (**Zhu et al. 2011**). The cellular and metabolic events that could influence the role of WT1 in the microRNA processing pathway also need to be investigated. It is quite possible that such tissue specific factors like WT1, would respond to different signals, as opposed to general transcription factors and ubiquitously expressed RNA binding proteins (**Treiber et al. 2017**). The microRNA pathway has been recently explored for early diagnostic purposes as well as therapeutic targeting. Circulating microRNAs can be detected from body fluids at very initial stages of diseases and thus help with early detection (**Wang et al. 2018**). However, the most promising therapeutic strategy seems to be the IGF signalling pathway. Incidentally, IGF1R has been tested as a therapeutic target in several cancers, including cellular models of WT (**Crudden et al. 2015, Bielen et al. 2012**). Our study not only provides a mechanistic insight into the causes of WT but also suggests potential diagnostic and therapeutic strategies involving the microRNA let-7 and the IGF signalling pathway.

## Materials and Methods

### FLASH and RIPseq

were performed as described earlier in **Bharathavikru et al. 2017**. Briefly, cells were crosslinked either by UV followed by formaldehyde or formaldehyde alone and subjected to Immunopulldown of WT1-RNA complexes through an antibody against WT1. The immune complexes thus precipitated where subjected to adaptor ligation and sequencing to identify the RNA subtypes that are associated with WT1.

### microRNA purification

Total RNA from cell lines was isolated as per the manufacturers’ protocols using the Qiagen RNeasy kit. Total RNA from kidney mesenchyme was purified using TRIZOL and ethanol precipitation. RNA was quantified using nanodrop and agilent bioanalyzer to verify the concentration as well as the purity of isolated RNA. WT patient FFPE samples were processed for xylene deparaffinization, followed by microRNA purification following the manufacturers’ protocol in the Qiagen FFPE miRNeasy kit.

### microRNA northern blotting protocol

microRNA was either selectively purified or total mRNA was used with probes specific for microRNAs to facilitate detection. DNA oligo was designed by using the template for antisense probe, synthesized oligo was resuspended in nuclease free water so as to obtain a 100μM stock solution. This oligo template was hybridized to T7 promoter primer and heated to 70 degC for 5 mins followed by room temperature incubation for 5 minutes to facilitate hybridization. Hybridized oligo was processed for klenow filling using exo-klenow, incubated at 37 degC for 30 mins followed by the transcription reaction catalysed by T7 RNA polymerase in the presence of labelled UTP. For the process of northern blotting, 10% Urea-PAGE was used for electrophoresing 10μg of total RNA samples, were blotted onto Nylon membrane for 1 hour using TBE as the buffer. After transfer, the membrane was UV crosslinked and processed for prehybridization for atleast 4 hours at 40oC in 10 ml of hybridization buffer. Following prehybridization, the labeled probe was added. After an overnight hybridization at 40⁰C, the samples were washed with 50ml of pre-warmed wash buffer followed by exposure to phosphorimager and an autoradiogram. Quantification of signals was performed using ImageJ analysis software, wherein U6 RNA levels were used as the loading control to normalize and calculate the expression difference across the test samples.

### Small RNA sequencing

Total RNA equivalent to 1μg was ligated with the 3’ RNA adapter with the help of the T4 RNA ligase 2 (deletion mutant) followed by ligation of the 5’ RNA adapter using T4 RNA ligase. The adapter ligated RNA was then subjected to reverse transcription with Superscript II RT followed by PCR amplification. The amplified cDNA was electrophoresed on a 6% PAGE gel. Mature microRNA, which is approximately 22 bp, gives an adapter ligated fragment of approximately 147 bp. A second band of 157 bp corresponding to piRNAs, other small noncoding RNA arising from 30 bp small RNAs was also excised together to give rise to the pool of small RNA. Additionally, the pre-processed forms of microRNA that result in longer fragments of approximately 160 bp to 300 bp were excised together for the pool of pre-processed small RNAs. The gel purified cDNA libraries were validated and sequenced on an Illumina platform.

### Dox inducible expression

Mcherry tagged Wt1 construct with doxycycline inducible CAG promoter was subjected to transient transfection in *Wt1* KO embryonic stem cells. Upon dox induction using 1μg/ml doxycycline, cells were processed for imaging, western blotting or RNA isolation and cDNA synthesis as described.

### qPCR for detection of microRNA

Pre-mature microRNA was quantified by designing primers against the full length of the premature microRNA. cDNA synthesis was carried out using total RNA and random primers. This was then processed for qPCR analysis using SYBR green based detection methods. Fold change was calculated by normalizing to U6 snRNA or 18S rRNA and then comparing the control (atleast n=2) to the mutant lines (atleast n=3). For mature microRNA expression, the clontech mir-X microRNA first strand synthesis and SYBR kit was used which involves an initial polyadenylation reaction and synthesis. MicroRNA detection was carried out by amplifying the cDNA with a 3’ primer that recognizes the polyadenylated fragments and the specificity is brought about by a 5’ microRNA specific primer (that corresponds to the entire mature microRNA sequence) and SYBR green based detection. Calculations were carried out as explained above. This methodology was used for detection of mature microRNA in the FACS sorted kidney mesenchyme tissue as well as the microRNA from the FFPE tissue.

### RNA Immunopulldown (RIP)

RIP was performed as described (**Bharathavikru et al. 2017**). Briefly, cells were formaldehyde crosslinked, and sonicated, followed by immuno pulldown with WT1 antibody. After overnight incubation and washes, samples were reverse cross linked, proteinase K treated and RNA was precipitated. Primer pairs specific for pre microRNAs and mature microRNAs were used with the clontech mir-X microRNA first strand synthesis and SYBR kit for detection of immune precipitated microRNAs and Northern Blotting as described.

### Protein immunopulldowns (IPs)

Protein IP was performed as described in Bharathavikru et al., 2016. Briefly, cells were lysed in RIPA buffer and IPs with WT1 and control IgG antibodies was performed in the presence of RNase. Resulting immune complexes were then resolved on 10% SDS PAGE gels and processed for immunoblotting.

### Immunoblotting analysis

Protein lysates and immune complexes from IPs were electrophoresed on 10% SDS PAGE gels and probed with antibodies against WT1 (ab89901), AGO1 (CST 9388), AGO2 (CST C34C6), DICER (CST D38E7), GFP (ab6556), LIN28B (CST S7056), DROSHA (CST D28B1), DDX5 (CST 9877), α/□-Tubulin (CST 2148S) Conformation specific secondary antibody (NEB L27AS).

### Gene ontology and Proteomics Analysis

A list of genes identified through Mass-Spectrometry analysis was selected for Gene Ontology (GO) analysis. The software tools Gorilla and ShinyGO 0.77 were utilized to analyze biological processes and construct a tree network of these processes. REVIGO was then used to highlight specific processes and visualize them in a scatter plot. To examine protein-protein interactions (PPI) and identify central interaction nodes, a STRING database analysis was performed.

### Chemically Induced Dimerization (CID) and FRET by Flow Cytometry

Detailed experimental protocol is provided in the Supporting Information. Briefly, the DDX5 CFP-FKBP and WT1-YFP-FRB vectors were created by routine cloning procedure. Following sequence confirmation of the clones, the vectors were subjected to a transient transfection in HEK293T cells. CID was induced by addition of Rapamycin and the resulting FRET change was measured using the Cytoflex flow-cytometer.

### DDX5 Knockout by CRISPR-Cas9 genome editing

Detailed experimental protocol is provided in the Supporting information. Briefly, the DDX5 knockout was performed by using a lentiviral CRISPR-Cas9 genome editing process. The knockout cell lines were created in the mesonephric M15 cells and validated using western blotting methods

### microRNA let-7 inhibitor experiment

M15, ML and MW cells were seeded in a 6 well plate and cultured for 24 hours. Cells that were 60% confluent were treated with 60nM of let-7 inhibitor (Ambion, Thermo Fisher Scientific Catalog No.-4464080) was transfected with FuGENE transfection reagent. Following 48 hours of incubation, the cell pellet was collected for RNA Isolation. cDNA synthesized from the isolated RNA and qPCR analysis to assess *Igf1r* expression was performed.

### Statistical Analysis

All experiments were performed with at least 3 biological replicates and technical replicates of each. Standard deviation was calculated to compute the error bars. Unpaired t-test was done to obtain the two-tailed p values, in graphpad prism. P value with significance levels are presented in the following way, ❬ 0.05=*, ❬ 0.01 =*\*\**, ❬0.001 =***, and ❬0.0001=****).

## Acknowledgments

The authors are grateful to Dr. Sara Macias Ribela, UoE (microRNA experimental advice), Dr.Cathy Duff (maintenance of transgenic mouse lines), Elizabeth Freyer (FACS facility), iGEM 2020 team for plasmids, Sequencing Facility, ILS, Bhubaneswar and CAIF facility IISER Berhampur.

## Author contributions

RSB, NDH, designed the study, SD, IJ, UB, RSB, JC, JS, SA, analyzed data, SD, UB, RSB, JS, JC, GP, VS, performed experiments, RSB, JC, KPJ, DM, AE, NDH, provided ideas and suggestions, RSB, KPJ, NDH, supervised experiments, RSB, NDH, obtained funding, SD, RSB, JC, NDH, wrote the manuscript with inputs from all authors.

## Data and materials availability

GEO accession id GSE169684 - MicroRNA profiling of Wilms Tumour 1 deleted and induced embryonic stem cells.

